# Ancient persistence and newfound diversity of CR1-group retrotransposons across chordates

**DOI:** 10.64898/2026.01.13.699120

**Authors:** Alexander J. Stuart, Zhenglong Du, Nozhat T. Hassan, David L. Adelson

## Abstract

Retrotransposons are mobile, repetitive DNA sequences that are ubiquitous across eukaryotes and widely recognised as key drivers of both gene and genome evolution. The CR1 group of retrotransposons is thought to have been present in the most recent common ancestor of chordates ∼560 mya, and is the dominant retrotransposon in the majority of chordate species. The advent of long-read sequencing technologies has enabled the assembly of high-quality genomes from representatives of almost all major chordate orders, enabling comparative analysis with deeply divergent species. To better understand the composition of CR1-group elements (CGEs) in chordates, we systematically characterised full-length, recently active transposable elements across representative species from every available extant order of chordate. Our analysis uncovered previously unknown phylogenetic relationships of CGEs within and between species and has pushed back the origin of certain CR1-group subclades by tens of millions of years. Additionally, entirely novel elements with no close relatives in existing databases were uncovered within several of the species analysed. We also detected numerous putative horizontal transfer events, many of which had not been previously documented. Overall, this investigation has provided the first chordate-wide analysis of an element that is historically understudied yet plays a pivotal role in genome biology and evolution.

## Introduction

Eukaryotic genomes are predominantly composed of repetitive sequences. These repeat-rich regions are known to be largely derived from Transposable Elements (TEs), sequences capable of moving and multiplying throughout the genome (Doolittle & Sapienza 1980). Their activity is known to influence the structure and size of genomes (Gable et al. 2024; Piskurek & Jackson 2012) and modify the function of genes both directly (Joly-Lopez & Bureau 2018) and through their regulation (Lynch et al. 2011; Sundaram et al. 2014), making them powerful drivers of evolution. The landscape of eukaryotic TEs is incredibly diverse, with many TE families originating early in eukaryotic evolution and subsequently diversifying within their host lineages (Smit 1996; Malik et al. 1999). In eukaryotes, the most abundant and active lineages of TE are the retrotransposons, elements capable of “copying and pasting” themselves into new genomic regions. Among these, Long INterspersed Elements (LINEs) are the most abundant in animals.

### Structural and evolutionary heterogeneity of LINEs

LINEs are defined by a highly conserved protein, ORF2p, containing both reverse transcriptase (RVT) and endonuclease domains that are essential for target-primed reverse transcription (Luan et al. 1993). While some LINE superfamilies require only ORF2p to be retrotranspositionally competent, the majority of LINE groups encode a smaller protein (ORF1p) towards the 5’ end. Initial ORF1 classifications recognised five types based on domain composition (Khazina & Weichenrieder 2009), but subsequent analyses have revealed a much broader and more mosaic diversity (Ivancevic et al. 2016). While the RVT domain is presumed to share a single ancestral source, ORF1 domains appear to be modular and frequently exchanged among LINE lineages and hosts (Metcalfe & Casane 2014). The function of ORF1 likewise varies between LINEs: in human L1 LINEs, ORF1 encodes the *transposase_22* (PF14244) domain, which acts as a nucleic acid chaperone, and is essential for retrotransposition (Cook et al. 2015); in some CR1 and L2 elements, ORF1 encodes an esterase-like protein implicated in membrane localisation, though transposition itself appears to not depend on this activity (Kajikawa et al. 2012).

### Origin, group and superfamily distinction of CR1

Chicken Repeat 1 Group (CR1) retrotransposons are a major group of LINEs first identified more than four decades ago (Stumph et al. 1981). As with many founding TE families, their early characterisation was ambiguous; the initial CR1 sequence was described as a chicken-specific Short INterspersed Element (SINE) lineage (Olofsson & Bernardi 1983), before longer homologous sequences were recognised across distantly related taxa. As the TE field matured, CR1 was elevated first to the rank of *superfamily* (Malik et al. 1999), and subsequently to the broader *group* level (Eickbush & Malik 2002), reflecting an increasingly recognised widespread distribution across metazoans. Currently, the CR1 group includes eight superfamilies: CR1, L2, L2A, L2B, Rex1, Daphne, Crack, and Kiri, which are widely distributed among all major phyla of Metazoans (Kojima 2020). CGEs, either existing as retrotranspositionally active copies or extinct fragments, are found across all classes of chordate (Shedlock 2006), and can make up large proportions of host genomes.

### Horizontal transfer

Horizontal transfer of transposable elements (HTT) is the process by which transposable elements move from the genome of one organism to that of another without vertical inheritance from parent to offspring. (Keeling 2024). While early research found infrequent transfer events between eukaryotes (Ivancevic et al. 2013), the addition of new taxa and TE libraries to subsequent analyses has shown that the frequency of HTT varies widely between different TE superfamilies and host lineages. Some taxonomic groups, such as ray-finned fish (Actinopterygii) and butterflies (Lepidopterans) have been found to commonly undergo HTT (Zhang et al. 2020; Reiss et al. 2019). Certain TEs, such as Rex1s, Mariners and hATs, exhibit greater rates of success in HTT between species than others (Zhang et al. 2020).

### Significance and aims of this study

Despite their ubiquity, CR1 group elements remain incompletely classified and annotated within major TE databases (e.g., Repbase, Dfam), and underexplored with respect to their rate of HTT. Here, we address these gaps by conducting a comprehensive survey of CR1 group elements across diverse major chordate taxa. Together, this study establishes an updated evolutionary framework for CR1 group elements and confirms a widespread and long-standing presence across chordates, which in turn has contributed to genome evolution in this group.

## Results

### Composition of CGEs within chordates

Using a combination of manual and automated curation methods, we characterised full length CGEs in genome assemblies for 101 species of chordate, using chromosome-level assemblies from NCBI. In the case of the class Aves, representative species from three major clades were chosen: Palaognathae, Galloanserae, and Neoaves. All 7 suborders from within the order Squamata were used. Both species of monotreme for which a genome assembly exists were used. For all other genomes, representative species were selected from every order for which a chromosome-level assembly was available (Supplementary Table 1). The genome sizes of our selected species ranged from 400Mb to 40Gb, spanning two orders of magnitude.

Using our pipeline, 1003 full-length CGEs (Supplementary Table 2) were identified in 73 of the 101 species, with no full length CGEs found in any of the 24 therian mammal species, consistent with previously reported research. Additionally, no full-length CGEs were detected in three species of teleost, representing the Percopsiformes, Gonorynchiformes and Ophidiiformes Orders.

The genomic proportion covered by CGEs (Figure 1) varied widely among species, ranging from 1.5% coverage in *Gambusia affinis* (Mosquitofish) to 47.9% coverage in *Tachyglossus aculeatus* (Echidna). All other species investigated exhibited CGE coverage from between 1% and 16%, with the notable exception of the Elasmobranchii (sharks and rays), exhibiting between 22% and 34% coverage.

**Figure 1:**
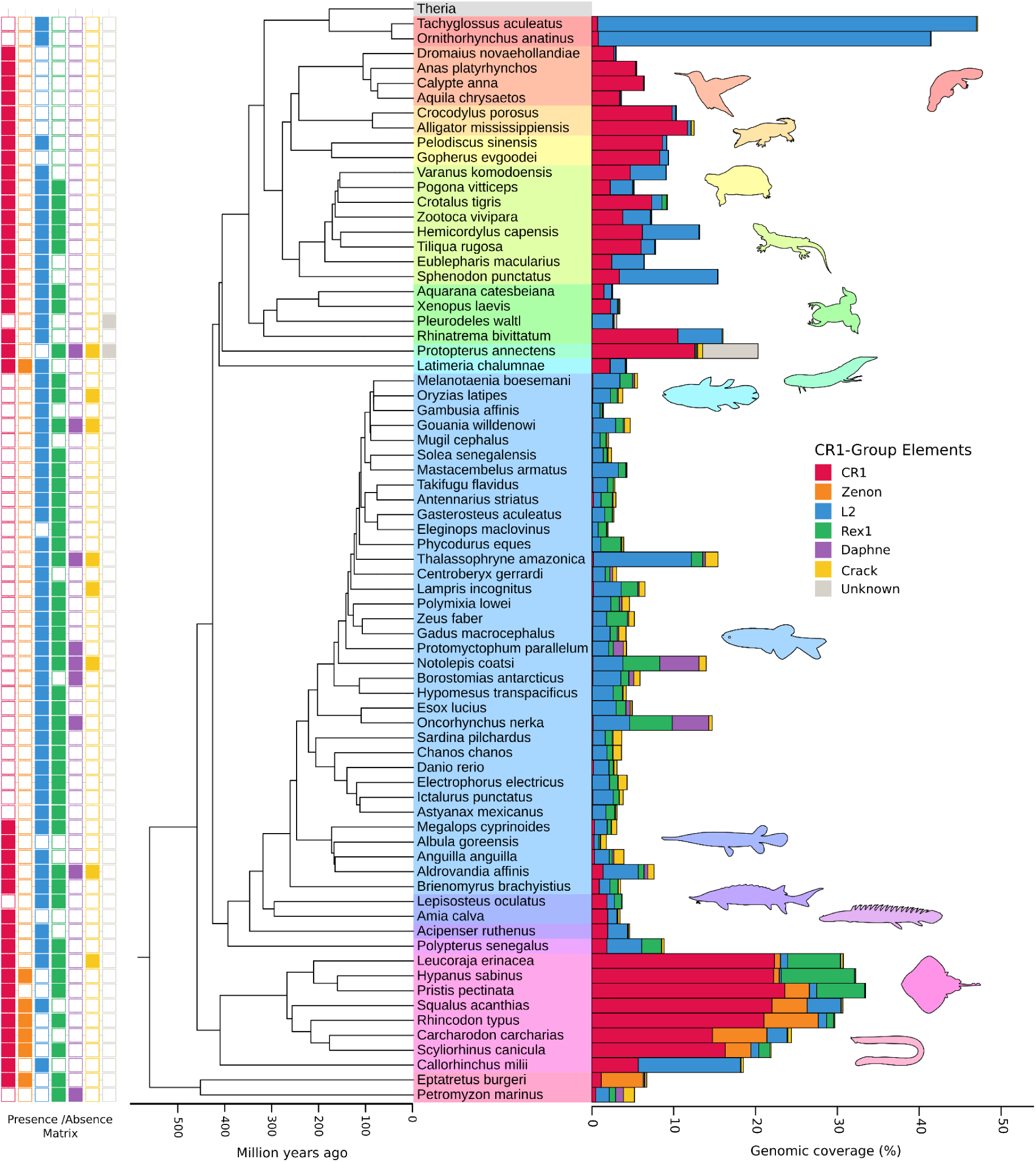
Phylogenetic relationships of the chordate species included in this study. A time-scaled tree was obtained using divergence estimates from TimeTree. The 24 therian mammal species were collapsed into a single grey node. Major clades are indicated by colour in order of appearance: red = Monotremata, orange = Aves, light orange = Crocodilia, yellow = Testudines, light green = Squamata, green = Amphibia, aqua = Dipnoi, light blue = Actinistia, blue = Teleostei, dark blue = Holostei, dark-purple = Acipenseriformes, light-purple = Polypteriformes, pink = Chondrichthyes, and light-pink = Agnatha. A stacked bar plot shows genome coverage (%) by CGEs in each species, with colours indicating the relative contribution of each clade. A presence-absence matrix of CGE clades having full length representatives is shown to the left of phylogeny.

Rex1 elements were present at low but persistent levels across most species, with mammals, birds, crocodilians, and turtles exhibiting particularly low coverage, all below 0.02%; Supplementary Table 3). The species with the highest Rex1 coverage were the Batomorphi (rays), *Oncorhynchus nerka* (salmon) and *Notolepis coatsi* (barracudina).

Across the sampled taxa, full-length CR1s were absent from five distinct chordate groups: mammals, Elopocephalai, *Lepisosteus oculatus* (spotted gar), *Pleurodeles waltl* (Iberian ribbed newt) and *Petromyzon marinus* (sea lamprey). Elasmobranchii (sharks and rays), coelacanth and hagfish were the only chordates found to have full-length Zenon elements. Full-length L2 elements, while not present in therian mammals, were detected in the majority of sampled chordates, including 55 species, and were especially prevalent among teleosts, where only five of the 38 genomes lacked detectable full length copies. Rex1 elements showed a more heterogeneous distribution across the species sampled.

Overall, the presence of Crack and Daphne elements did not follow a clear pattern. Full length elements are completely absent across Sarcopterygii, and only present in 10 teleost species, *Leucoraja erinacea* (little skate), and *P. marinus*. Notably, only 2 teleost species investigated appear to have had a major expansion of Crack elements, the same two species with the highest Rex1 coverage, *O. nerka* and *N. coatsi*.

### Broad-scale phylogenetic relationships between CGEs

The conserved amino acid RVT domain from both our *de novo* identified CR1 group elements and previously characterised CGEs obtained from Repbase and Dfam were used to infer a Bayesian phylogeny using BEAST (Supplementary Figure 1A). This analysis recovered four well-supported CGE clades, each receiving posterior probabilities between 0.92 and 1.00. In contrast, the deeper relationships between these clades were poorly resolved. The nodes forming the backbone of the tree generally exhibited low posterior probabilities (below 0.55), with the highest reaching 0.67. Although this value exceeds that of the other deep nodes, it remains far below the support values observed in the four well-supported clades, and remains insufficient to infer a reliable and high confidence branching order among the clades.

Classification of Repbase and Dfam elements within these clades showed several inconsistencies, with a small number of TEs assigned to different superfamilies than the majority of sequences in their respective clade. To address this, we designated clades containing representatives from different CGE superfamilies as “Canonical” when the original, first-described sequence from that superfamily was present in that cluster. Three clades contained over 100 members and were designated as the Rex1, Canonical-CR1/Zenon and L2/Daphne/Crack clades. Additionally a clade comprising four *de novo* salamander-derived sequences was recovered. The Rex1, Canonical-CR1/Zenon and L2/Daphne/Crack clades of CGEs included elements originating from a wide range of Metazoan phyla. The phylogenetic diversity of the host species, coupled with the high clade confidence values, provides strong evidence for the monophyly of these four groups, and suggests an ancient origin for each of these major clades.

Further investigation of the L2/Daphne/Crack clade showed that it forms a low support polytomy comprising seven high-confidence subclades (Supplementary Figure 1B). The largest of these subclades contained 168 sequences, all of which were classified as L2 or Kiri elements. The second largest clade, with 73 members, was primarily composed of Daphne elements, along with a small number of Repbase derived sequences classified as CR1s. Crack elements were distributed across three distinct subclades. The largest of these contained 44 sequences classified as Crack or L2B from a wide range of phyla, including chordates. Two additional Crack subclades, comprising 13 and 7 sequences respectively, were composed exclusively of chordate-derived elements. Two sequences classified as L2A formed a distinct, well-supported clade. Finally, two sequences characterised from the *Protopterus annectens* (lungfish) genome formed a distinct subclade and could not be confidently assigned to a superfamily.

### Clade level phylogenetic structure of CGEs

Using maximum likelihood methods, we reconstructed the phylogenetic relationships between Canonical-CR1/Zenon and Canonical-L2 CGEs. For these analyses, we used consensus sequences generated from chordate TEs characterised in this study that were represented in the genome by a sufficient number of copies exceeding a minimum length, enough to reliably represent the ancestral elements that gave rise to extant genomic copies. This was combined with sequences from Repbase and Dfam, to help orient the position of chordate CGEs within the greater Animalia phylogeny.

#### The Canonical-CR1/Zenon Clade

The Canonical-CR1/Zenon clade (Figure 2) contained 204 sequences from a diverse range of eukaryotic phyla. All chordate CR1s were recovered in a single well-supported clade containing 49 consensus sequences, composed of 48 from chordate genomes (with five previously catalogued in Repbase and Dfam) and one Repbase derived sequence from *Chionoecetes opilio* (snow crab) with 97% bootstrap support. Within this clade, Avian, Crocodilian, Chondrichthyan and Actinopterygian sequences formed strongly supported monophyletic groups. Squamate CR1s were recovered in two distinct, well-supported clades, while turtle-derived sequences were distributed across three separate clades within the tree. We found that the snow crab CR1 had the highest amino acid sequence similarity in the RVT domain to a CR1 from *Callorhinchus milii* (Elephant shark) at 57%. Sequence homology of 57% is significantly below our threshold for HTT candidates (see below); however, we cannot exclude the possibility of an ancient HTT event.

**Figure 2:**
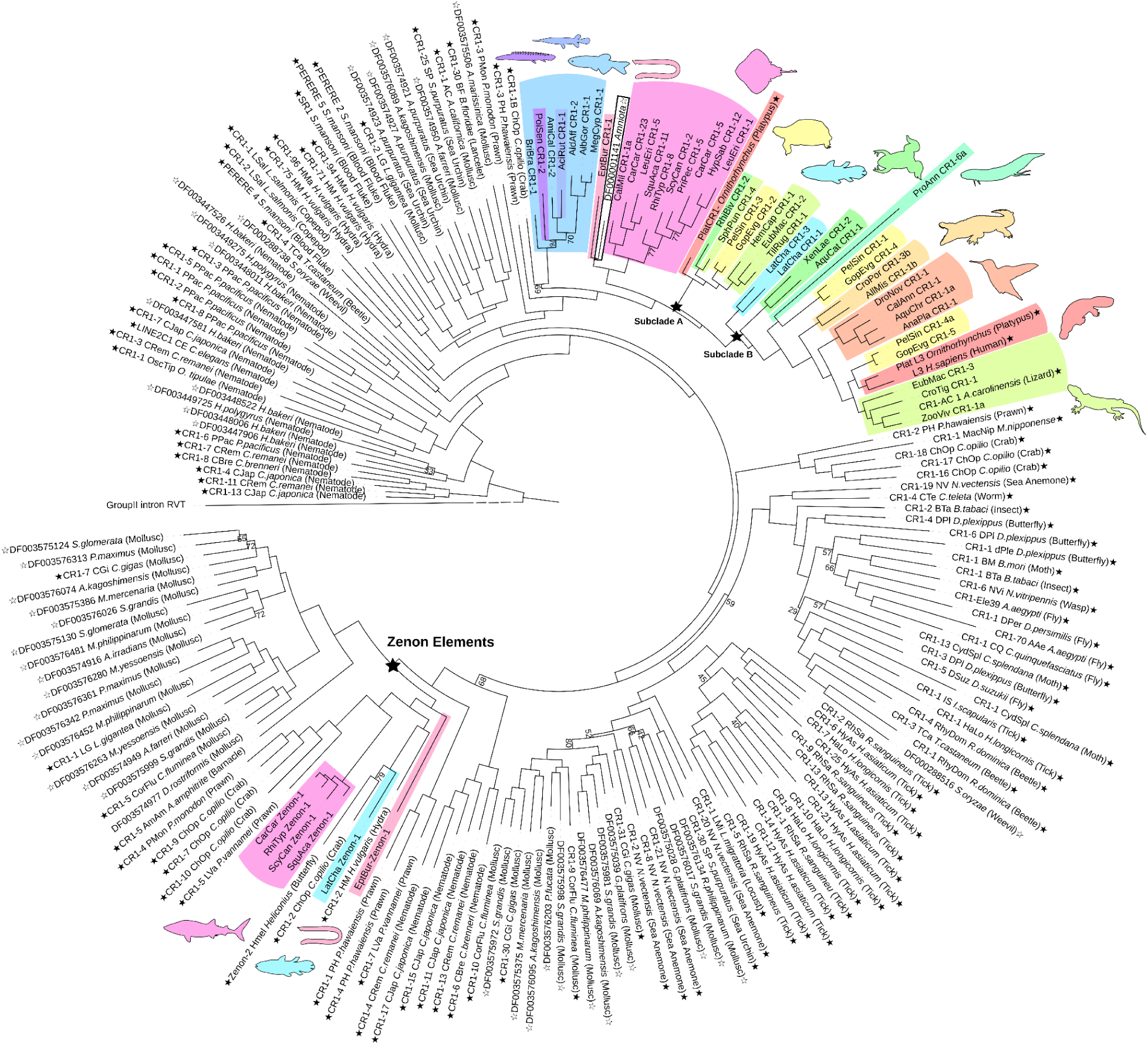
Phylogenetic relationships of the Canonical-CR1/Zenon Clade reconstructed with IQ-TREE2 from multiple sequence alignment of the conserved RVT domain. Major chordate lineages are indicated by colour, corresponding to major taxonomic groups. The Group II Intron is used as an outgroup (not to scale). Colours: purple = Polypteriformes, blue = Actinopterygii, dark blue = Acipenseriformes, light-pink = Agnatha, pink = Chondrichthyes, red = Mammalia, dark-green = Amphibia, yellow = Testudines, light-green = Lepidosauria, light-blue = Actinistia, aqua = Dipnoi, light-orange = Crocodilia, dark-orange = Aves. Bootstrap values below 80 are shown. RepBase and Dfam derived sequences are denoted with a black and white star respectively.

Full length Zenon elements were recovered from Elasmobranchii (sharks and rays), coelacanth and hagfish (Figure 2). Sequences were recovered from five Elasmobranchii species, with all full length consensus sequences exhibiting greater than 84% sequence identity to one another. Unlike CR1s and L2s, chordate Zenon elements did not cluster monophyletically but instead grouped with elements from disparate taxa (hagfish as the outgroup, coelacanth and elasmobranchs with different crustacean elements), with the reverse transcriptase (RVT) domain exhibiting low sequence identity across these pairings. The incongruence between the host phylogeny and the Zenon tree makes it difficult to infer the evolutionary history of these elements. Additional sampling from non-chordate taxa may improve our understanding of the relationships of Zenon elements between species. In the case of the hagfish, Zenon elements were the most abundant CGE, although in other genomes this was not the case.

#### The Canonical-L2 Clade

The Canonical-L2 clade comprised 169 consensus sequences originating from 56 chordate orders, including 13 chordate sequences originating from RepBase and Dfam, as well as 33 sequences from a range of other eukaryotic phyla. Maximum-likelihood phylogenetic analyses recovered all chordate L2s within two major lineages, designated Clade A and Clade B (Figure 3). Clade A contained 4 well-supported subclades (1A to 4A). Subclade 1A included elements from 4 sarcopterygian (lobe-finned fish) orders. Subclade 2A encompassed a broad range of deeply divergent chordate taxa, including actinopterygians, chondrichthyans, *P. marinus* and a reconstructed Amniote sequence. Subclade 3A was exclusively composed of Teleost elements, while subclade 4A was composed of Actinopterygian sequences, along with sequences from *L. erinacea*.

**Figure 3:**
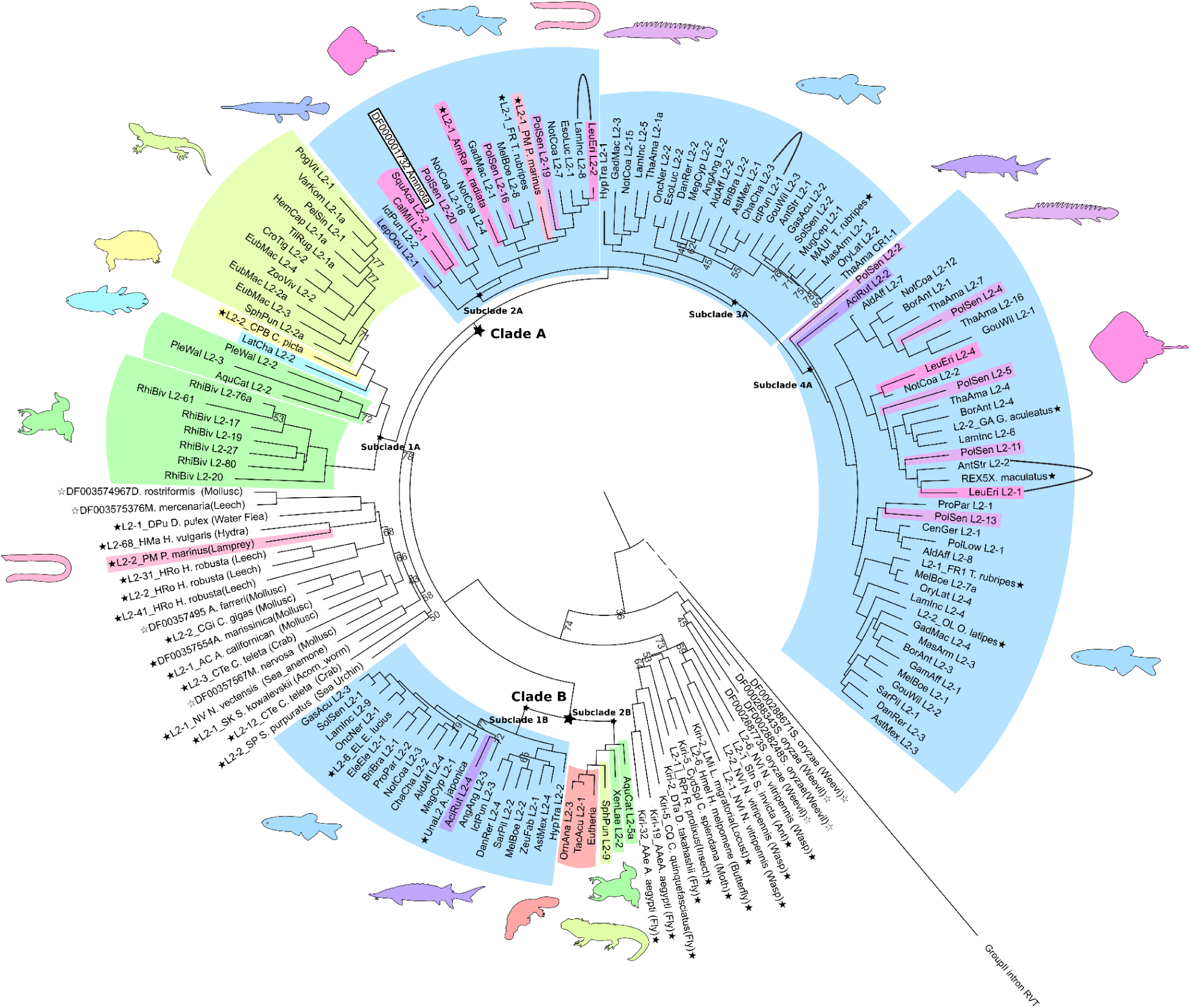
Maximum likelihood phylogeny of Canonical-L2 elements reconstructed with IQ-TREE2 from multiple sequence alignment of the conserved RVT domain. Major chordate lineages are indicated by colour, corresponding to major taxonomic groups. The Group II Intron is used as an outgroup (not to scale). Colours: light-pink = Agnatha, dark-green = Amphibia, light-blue = Actinistia, yellow = Testudines, light-green = Lepidosauria, blue = Actinopterygii, dark-blue = Holostei, pink = Chondrichthyes, dark-purple = Acipenseriformes, light-purple = Polypteriformes, red = Mammalia. Bootstrap values below 80 are shown. RepBase and Dfam derived sequences are denoted with a black and white star respectively. Putative horizontal transfers are marked with a black arc.

Clade B, separated into subclades 1B and 2B, forms a distinct branch, separated by sequences from a diverse range of non-chordate taxa. Relative to Clade A, Clade B was recovered in a smaller subset of chordate taxa. Subclade 1B contained sequences exclusively from teleosts and *Acipenser ruthensis* (sturgeon), while subclade 2B contained sequences from mammals, anurans and *Sphenodon punctata* (tuatara).

The Repbase-derived *P. marinus* (Lamprey) sequence L2-2_PM did not cluster within either major clade but instead grouped most closely with L2-68_Hma, another RepBase sequence originating from *Hydra*. Despite this placement, the two sequences shared only 47% amino acid identity across the RVT domain. Additional screening detected no copies of this *P. marinus* repeat in its closest sampled relative, *Eptatretus cirrhatus* (hagfish), suggesting that this L2 is not broadly distributed across Agnathans. In the absence of any closely related elements, the evolutionary relationships of this L2 to other CGEs remains unresolved.

Canonical-L2 retrotransposons were detected in the majority of chordate lineages examined, with only 46 of the 101 species lacking multiple full-length copies (Figure 1). Among these, six lineages – therian mammals, birds, *Gopherus evgoodei* (thornscrub tortoise), *Protopterus annectens* (lungfish), *Amia calva* (bowfin), and Galeomorph sharks) – lack L2s. In each of these groups, the closest relatives that retain L2s diverged more than 150 million years ago, suggesting that the losses are ancient and independent. In contrast, four teleost species exhibit L2 losses despite the presence of active elements in closely related taxa. Overall, these patterns suggest that L2 elements were likely present in the last common ancestor of chordates, and that their loss has been infrequent at deep phylogenetic levels, occurring only sporadically in individual lineages.

While the dynamics of Sarcopterygian L2s appear to generally align with the species phylogenies for L2s, with the exception of the placement of LatCha_L2-2 in Subclade 1A (Figure 3), the dynamics of the Actinopterygian L2s are more difficult to understand. The inclusion of the Chondrichthyan and lamprey sequences within otherwise wholly Actinopterygian monophyly are unexpected, suggesting possible ancient HTT events.

### General structure of chordate CGEs

Across chordate CGEs, the ORF2 displayed typical canonical LINE domain organisation, consistently containing both an apurinic endonuclease (APE) and RVT domain. ORF1 showed heterogeneous presence and domain architecture (Figure 4). Rex1 and Zenon elements universally lacked ORF1 in all surveyed sequences. By contrast, esterase-type ORF1s were the only identifiable ORF1 subtype in CR1 and L2 lineages recovered.

**Figure 4:**
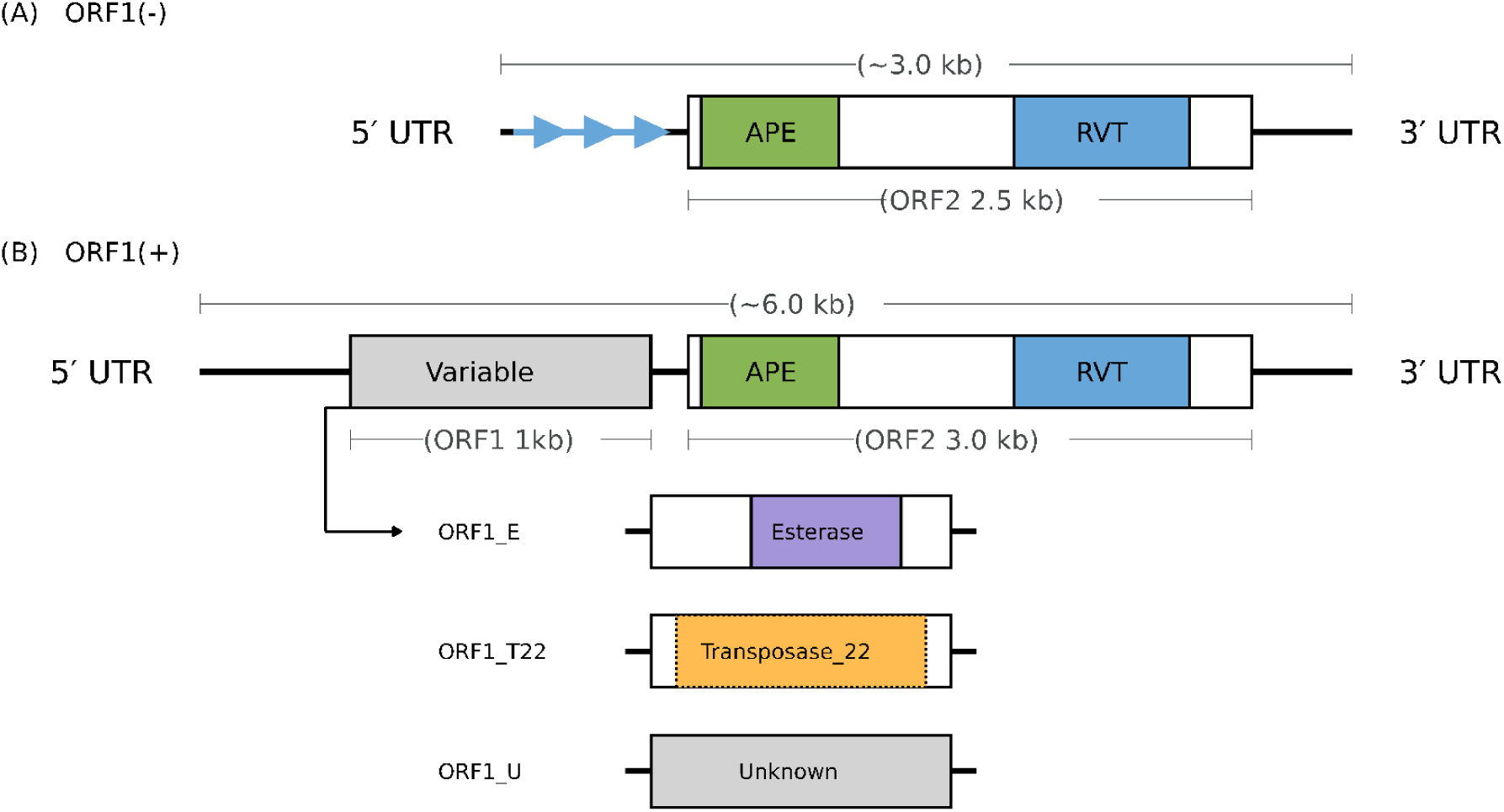
Structural variants of chordate CR1 Group Elements. CR1 Group elements display two principal structural organisations. **(A)** The shorter variant encodes a single open reading frame (ORF) corresponding to the reverse transcriptase (RVT), here designated ORF1(–). **(B)** The longer variant encodes an additional upstream ORF, designated ORF1(+). In some ORF1(–) elements, a repeating subunit (1-5 subunits per array with subunit lengths and sequences species specific, ranging from 100 to 500bp) occurs near the 5′ UTR. APE; Apurinic/apyrimidinic endonuclease. RVT: reverse transcriptase domain.

All chordate Canonical-CR1s (Figure 2) either encode an esterase ORF1 or cluster with esterase-encoding relatives that do, suggesting convergent secondary ORF1 losses rather than independent gains of an esterase ORF1. Zenon elements included in this study from all phyla exhibited two consistent features, in that they lacked an ORF1 and possessed a poly-A tail. Among the CGEs, the presence of a poly-A tail is unique to Zenon elements.

Within L2 elements, two major clades are present in chordates (Figure 3). Clade B lacks ORF1 entirely, whereas Clade A shows variable ORF1 retention. Most Teleostei and Polypteriformes encode esterase ORF1s, while Holostei, Acipenseriformes, and Chondrichthyes lack ORF1 but cluster with esterase-encoding lineages. Lepidosauria and Coelacanthiformes lack ORF1, and Amphibia contain predominantly ORF1(-) L2s, with only rare representatives clustering among ORF1(+) lineages (Supplementary Figure 2).

Inclusion of additional lineages currently assigned to the CR1 group reveals broader diversity. Daphne elements form a monophyletic clade within the broader L2/Crack/Daphne group (Supplementary Figure 1B). Chordate derived Daphne ORF1s exhibit homology to transposase-22 but with low sequence similarity (Supplementary Figure 3), whereas no ORF1 could be reliably detected for chordate derived Crack elements.

Overall, ORF1 in CGEs show recurrent loss and replacement across deep evolutionary time. ORF1 presence/absence and domain identity do not map cleanly onto current superfamily boundaries, indicating frequent secondary loss and modular domain turnover rather than a conserved ORF1 architecture within the CR1 group.

### CGE clades of unresolved affiliation

The genomes of two species, the salamander *Pleurodeles waltl* and the lungfish *Protopterus annectens*, contained TEs with poorly resolved affiliations (Supplementary Figure 1). The *P. waltl* sequences did not cluster with any other CGEs in the Bayesian tree, while the *P. annectens* sequences were nested within the broader L2/Crack/Daphne clade. Pairwise alignment showed that these novel sequences are substantially divergent from all other characterised CGEs, with the conserved RVT domain exhibiting less than 50% amino acid identity to TEs identified within Repbase and Dfam as CR1s, Cracks and Daphnes. Six unclassified *P. waltl* TEs were characterised, making up ∼0.003% of the respective host’s genome, while 72 such elements were found in *P. annectens*, making up ∼6.9% of this genome. Additionally, it was found that these *P. waltl* TEs carry an ORF1 with no detectable similarity to known protein families.

### Horizontal transfer between chordates

Our methods for HTT inference were broadly similar to previous studies (Muller et al. 2024; Zhang et al. 2020), and were able to identify 95 candidate pairs (Figure 5; Supplementary Table 4). From our analysis, we observed that Rex1 elements are the most frequently horizontally transferred TE within the CR1 group. We also observed a few instances of L2 transfer, and only three CR1 transfers between species. Almost all the HTT events occurred between teleosts, with the highest number of putative transfers involving *A. affinis* (ray-finned fish). We also observed multiple putative HTT events involving a Rex1 element between *P. marinus* and several ray-finned fish.

**Figure 5:**
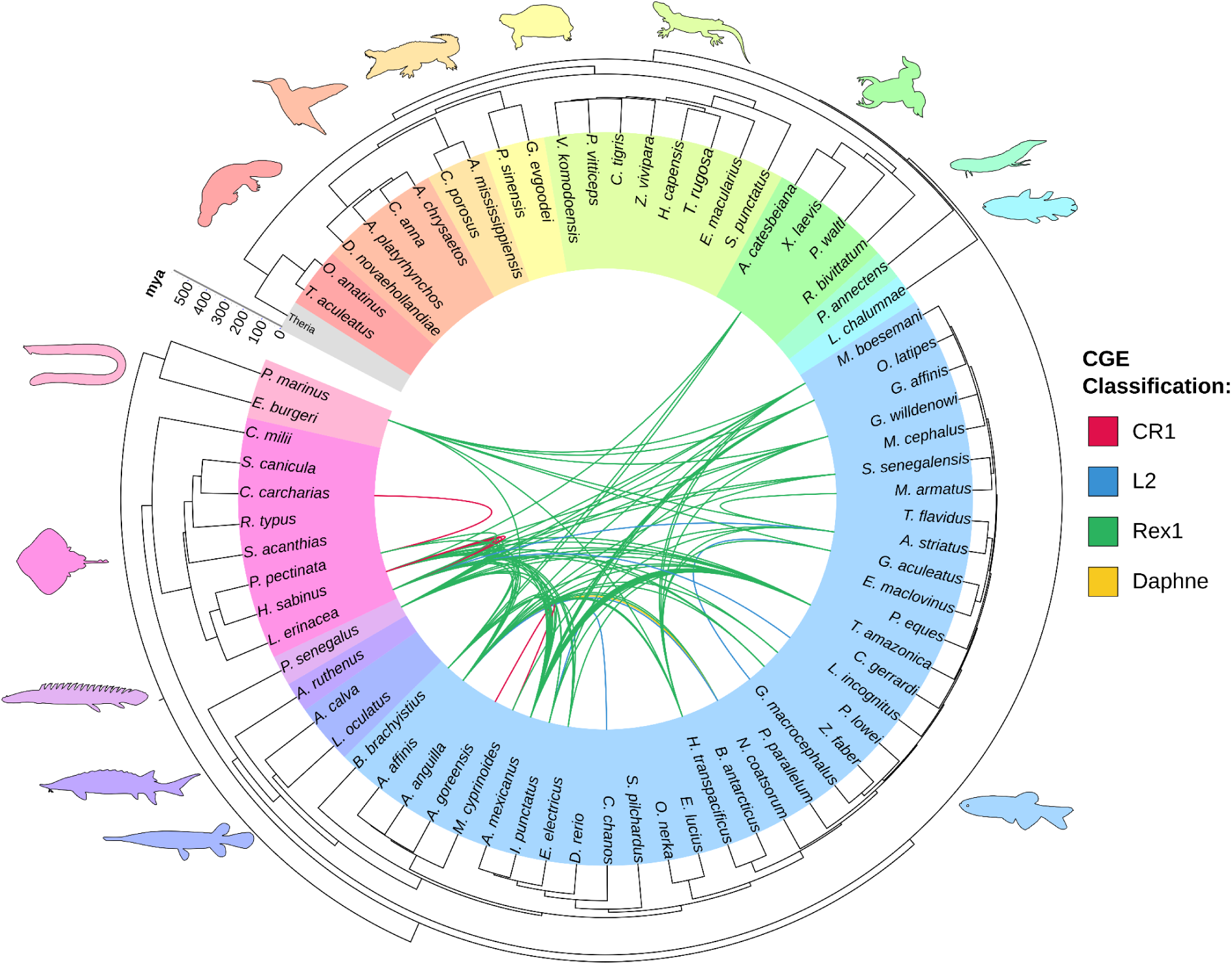
Inferred horizontal transfer events. Host phylogeny represents species analysed in this study with each major chordate lineages indicated by colour in order of appearance: red = Monotremata, orange = Aves, light orange = Crocodilia, yellow = Testudines, light green = Squamata, green = Amphibia, aqua = Dipnoi, light blue = Actinistia, blue = Teleostei, dark blue = Holostei, dark-purple = Acipenseriformes, light-purple = Polypteriformes, pink = Chondrichthyes, and light-pink = Agnatha. Each curve connecting two species in the host tree represents one of 95 inferred HTT events. The scale on the left hand side shows time in MYA. Curves indicate HTT of each CGE categorised by colour with red representing CR1, blue L2, green Rex1, and yellow Daphne.

### Re-evaluation of reference CGE classifications

To evaluate the fidelity of both sequences existing in databases and chordate CGEs characterised in this study, the sequences from the Bayesian BEAST tree (Supplementary Figure 1) were used to perform an all-versus-all pairwise similarity search (Figure 6). For each CGE, we compared its highest scoring similarity hit with its placement in the phylogeny to identify cases where annotations conflicted with phylogenetic relationships. As expected, percent sequence similarity negatively correlates with patristic distance, with increasing spread at the lower end of similarity. In our dataset, we found the highest percent similarity at which a closest pairwise match was observed between two sequences from different superfamilies, both in their BEAST clade assignment and TE classification, was at 48% similarity. Strikingly, there is an alarming proportion of database annotations that conflict with our phylogenetically defined groupings, even well above the twilight zone of protein sequence similarity (Rost 1999; Chang et al. 2008).

**Figure 6:**
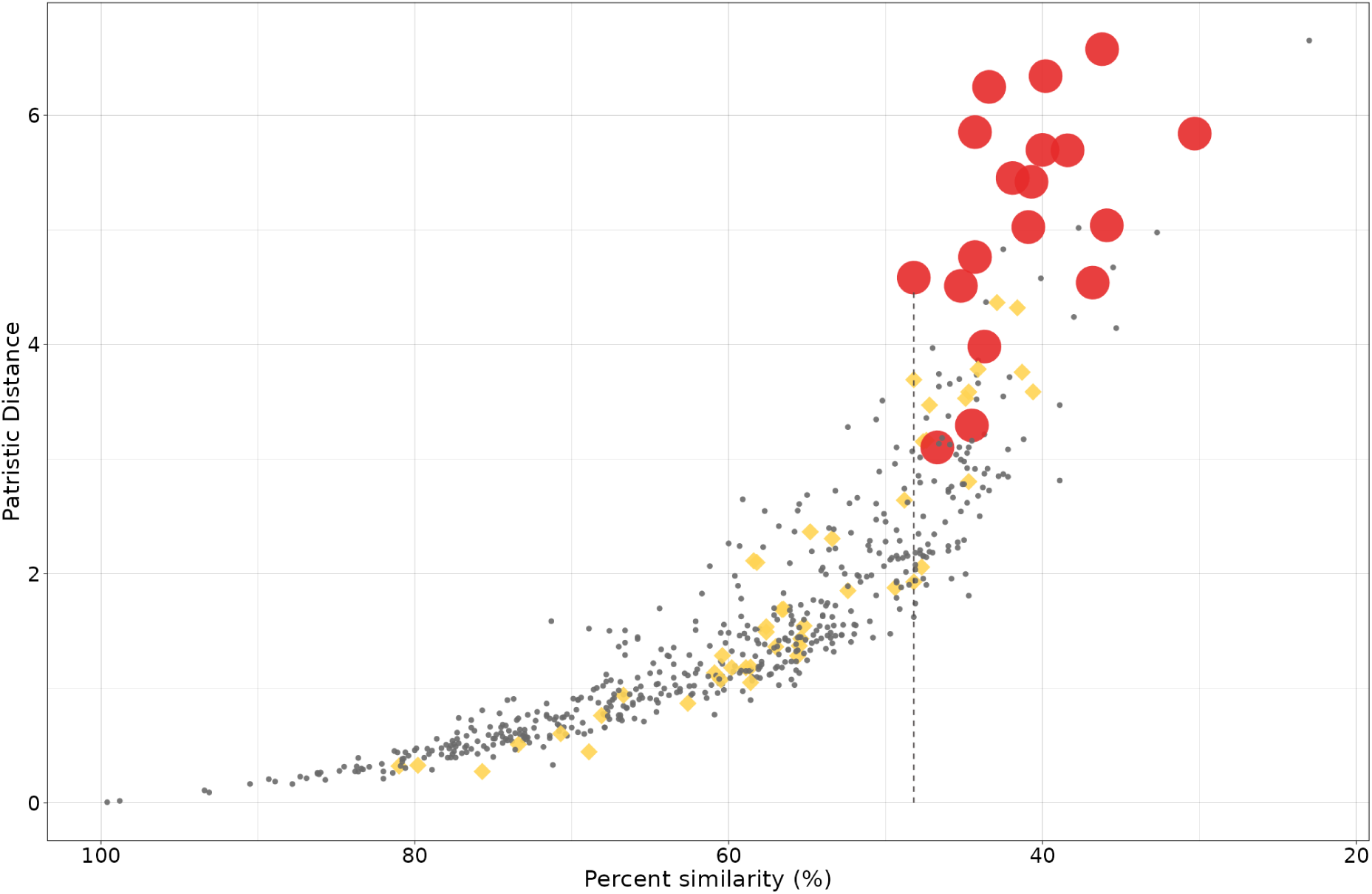
Pairwise similarity profiles of novel and previously characterised CR1-group elements used in phylogenetic reconstruction. An all-versus-all pairwise alignment was performed for elements included in the BEAST RVT tree (Supplementary Figure 1), with the top match for each sequence recorded. For each highest scoring hit, percent similarity is compared to patristic phylogenetic distance. Yellow diamonds indicate database annotations that conflict with phylogenetic placement, that is when the pairs are assigned different superfamily classifications despite occurring in the same reconstructed clade, or are assigned the same superfamily but placed in different clades. Large red dots represent pairings that do not occur in the same reconstructed phylogenetic clade. The dashed line denotes the highest percent identity value of a pair belonging to separate clades.

## Discussion

### Horizontal transfer

We will discuss the results of the horizontal transfer analysis first, as they are relevant to our later discussion of the phylogenetic analyses. Our HTT analysis suggests that some clades of the CR1 group appear to exhibit a high tendency for transfer, with others transferring rarely or not at all (Figure 5). Rex1 elements were the most frequently transferred CGE, with most events occurring between teleosts, supporting previous reports on Rex1 mobility and the observation that HTT is more likely to occur between species with close evolutionary relationships (Zhang et al. 2020; Muller et al. 2024; Ivancevic et al. 2013). In contrast, we observed limited HTT events involving Canonical-CR1 and Canonical-L2 elements, which is also supported by previous findings (Zhang et al. 2020; Volff et al. 2000). We also observed no HTT events within amniotes, which is unusual as HTT of LINEs has previously been documented in amniote species (Zhang et al. 2020; Muller et al. 2024; Walsh et al. 2013). Interestingly, a high number of Rex1 and Canonical-L2 HTT events were observed between Batomorphi (rays) and teleosts, despite the substantial divergence time of ∼462 mya between them (Kumar et al. 2022). No ORF1 has been identified in any Rex1 element, either in our dataset or in previously published work. This structural feature is shared with other TE clades known for frequent HTT (Walsh et al. 2013; Malik & Eickbush 1998). While we hypothesise that Rex1 may have structural characteristics contributing to their high mobility, what governs their increased rate of HTT over other CGE clades remains unknown.

### Phylogenetic structure and lineage loss in CR1-Group elements

#### Canonical-CR1 Elements

The Canonical-CR1/Zenon tree (Figure 2) recovered a smaller monophyletic group containing all chordate CR1s. The broad distribution of CR1s across Eukaryotic phyla is well established (Malik et al. 1999), and suggests the clade as a whole is either ancient in origin or the subject of frequent HTT. However, CR1s have been shown to be rarely horizontally transferred (Zhang et al. 2020; Peccoud et al. 2017), a pattern supported by our own HTT analysis. While previous analyses have suggested that CR1s were present in the ancestral amniote (Suh et al. 2015; Gable et al. 2024; Shedlock 2006), our tree reconstruction provides evidence for an even older, monophyletic origin of CR1s in the most recent common ancestor (MRCA) of chordates.

Prior work has suggested that the common ancestor of amniotes contained at least two distinct lineages of CR1 (Gable et al. 2024). With the inclusion of additional taxa in our analysis, we observed that CR1s from the more distantly related amphibians, lungfish and coelacanth did not form an outgroup but instead clustered into the two clades previously reported (annotated as subclade A and B, Figure 2). Although ancient HTT events could explain this pattern, an explanation based on vertical transmission is more parsimonious based on our observed rates of CR1 HTT in chordates. This in turn suggests that Subclade A was present in the MRCA of tetrapods at least ∼352mya, while Subclade B was present in the MRCA of sarcopterygians at least ∼415mya.

#### Zenon elements

Zenon elements are a poorly characterised clade of CGEs, first identified as a CR1 lineage in butterflies (Novikova et al. 2007), and subsequently identified and formally described in a limited number of taxa, including insects, crustaceans, molluscs and hydra. Despite the fact that no chordate TEs annotated as Zenon elements are present in existing repeat databases, our pipeline detected them in three chordate lineages: hagfish, coelacanth, and Elasmobranchii. The hagfish Zenon represented a novel TE, while the coelacanth Zenon is misannotated as CR1-5_LCh in Repbase. None of the elasmobranch Zenon elements have been formally annotated and submitted to a repeat repository, but were correctly identified in the whale shark genome assembly (Weber et al. 2020).

#### Canonical L2-Elements

The presence of multiple lineages of L2, corresponding to clades A and B, has been reported in datasets with limited chordate representation (Sugano et al. 2006; Gemmell et al. 2020). Our analysis shows that Clade B has a far more restricted distribution in chordates than the more common Clade A, being found in just over half of all Actinopterygian representatives, along with the monotremes, tuatara and anurans (Figure 3). In all species containing Clade B L2s, with the exception of the monotremes, Clade A L2s are also present. A defining feature of Clade B L2s characterised in this study is the absence of an esterase ORF1. Although this difference of esterase ORF1 presence within species was previously documented in zebrafish (Sugano et al. 2006), our findings indicate that it represents a broader distinguishing feature between Clade A and Clade B L2s across chordates.

### Broader species-level trends

Previous work has not shown a clear relationship between intraspecies TE diversity (the number of unique and deeply divergent TE lineages) and abundance (Elliott & Gregory 2015), a finding consistent with our own results. Here we look at CGE lineages, described as a species-specific cluster of CGEs sharing more than 60 percent identity across the RVT domain, a heuristic threshold chosen based on our pairwise similarity analyses (Figure 6). Using this definition, *T. aculeatus* shows the highest CGE abundance at 47.9 percent genome coverage but contains only a single CGE lineage, whereas the teleost *N. coatsi* exhibits 14.2 percent CGE coverage yet contains 24 CGE lineages (Supplementary Table 5). In our sampled chordate species, amniotes exhibited consistently low diversity, typically containing three or fewer CGE lineages. In contrast, teleost and Chondrichthyan species displayed a far greater range of CGE diversity, although no clear taxon-specific patterns were apparent. Characterisation of new teleost and Chondrichthyan CGEs in a broader range of species at a finer taxonomic level (e.g. the Class level) will be required to determine whether the observed variation reflects taxon-specific expansions or undescribed HTT events, both of which could increase CGE lineage diversity but on different evolutionary timescales.

Presence-absence analysis of recently active Canonical-CR1 group elements revealed a consistent pattern of retention across chordates. The current distribution of Canonical-CR1s can be explained with only five apparent extinction events, occurring in mammals, salamanders, Elopocephalai (a large teleost group comprising approximately 90% of all species, 31 out of 35 orders), bowfin and lamprey. These extinctions are inferred from the absence of recently active elements, as defined by our extraction pipeline criteria. We speculate that some form of selective pressure may act on these TEs, imposing an evolutionary constraint against their total extinction. A comparable pattern of long-term vertical retention and hypothesised selective pressure has been proposed in R2 LINE elements (Kojima & Fujiwara 2005; Kojima et al. 2016; Hassan et al. 2025), although no mechanism has been identified.

### Limitations of heuristic thresholds in TE annotation

A long-standing weakness in TE taxonomy lies in the use of heuristic criteria. A clear example is the pragmatic “80/80” (80% identity over 80% of sequence length) threshold, commonly used for defining TE families (Wicker et al. 2007). While widely adopted, this rule has been criticised for its arbitrariness and its failure to align consistently with evolutionary relationships (Piégu et al. 2015; Seberg & Petersen 2009). Within the CR1 group, individual clades were initially described based on phylogenetic reconstruction, but with sequences derived from a limited number of taxa (Eickbush & Malik 2002). As genome sampling has expanded, many additional sequences have been assigned to these clades solely based on pairwise sequence similarity. In practice, these assignments are often made because the existing clades provide the closest available match, but may not reflect true evolutionary relationships in more deeply divergent sequences.

Despite this, threshold-based annotation remains a necessary concession in classifying novel TEs, where heuristic criteria provide a pragmatic means of grouping related sequences. To assess the behaviour of such thresholds in practice, we compared pairwise similarity profiles with well-supported phylogenetic groupings. In our dataset, RVT-domain amino acid identities above 60% tended to align with phylogenetically defined groupings (Figure 6). Future classifications of novel TEs made based on an RVT percent identity match of less than 60% should be treated with caution, as similarity-based clustering below this threshold no longer corresponds reliably to phylogenetic structure.

### Re-evaluation of CR1 Group taxonomic boundaries

The instability of TE taxonomy is compounded by historical constraints. The initial CGE clades were defined when only a small number of genomes were available (Malik et al. 1999), and explicit annotation criteria were often lacking. As a result, public TE databases (e.g., Repbase, Dfam) contain numerous legacy annotations that do not reflect current phylogenetic knowledge. Because most annotation pipelines propagate these classifications, early misassignments continue to introduce systematic error into comparative and evolutionary studies (Hassan & Adelson 2023). Updating outdated and inaccurate resources with phylogenetically informed classifications is therefore essential.

In our analysis, we used two criteria to identify likely misannotations: (1) phylogenetic placement where a clear majority-minority pattern indicates incongruence, and (2) pairwise sequence identity (>∼60%) between elements assigned to different superfamilies. Misannotation under these conditions was common. A notable case involved teleost “CR1” elements, many of which cluster robustly within the Canonical-L2 clade and share high sequence similarity with known L2 elements. These appear to trace back to a misannotation of the MAUI element (Poulter et al. 1999) in Repbase, resulting in incorrect propagation of the CR1 label across teleost genomes. After reassigning these sequences to L2, no clear evidence remains for genuine, recently active Canonical-CR1 elements in non-Elopocephalai teleosts.

At broader scales, most CGEs maintain their place within the currently accepted taxonomy. Rex1 formed a distinct, monophyletic and well-supported lineage across analyses. Zenon elements, which are considered to be a subcategory of CR1s by Dfam, were found to robustly cluster with Canonical-CR1 elements in our Bayesian reconstruction. Despite this, Zenon retains a distinct molecular signature, most notably the presence of a poly-A tail, unique among CR1-group elements.

The relationships among the L2, Kiri, Daphne, Crack, L2A, and L2B superfamilies are more complex. Dfam currently subsumes all into L2, a choice that is broadly congruent with phylogeny but collapses substantial internal evolutionary and structural diversity. Conversely, the Repbase scheme (Bao et al. 2015) treats these as parallel superfamilies, which is not fully supported: Kiri was found to nest within the Canonical-L2 clade, while elements labelled as Crack form multiple subclades rather than stable independent lineages. By comparison, sequences classified as Daphne formed a strongly supported monophyletic group spanning multiple Metazoan phyla, indicating a shared evolutionary history and supporting its recognition as a distinct superfamily. Together, these patterns suggest that CR1-group boundaries, particularly within the L2-derived lineages, require refinement to reflect evolutionary history while preserving meaningful internal structure. Additional sampling from non-chordate species is needed to more clearly understand the relationships within CGE superfamilies in metazoans.

Additionally, the ancient origins of these clades make it likely that even the conserved ORF2 RVT domain has become saturated by multiple substitutions. It is likely that this saturation has contributed to our inability to resolve the higher level relationships in the BEAST topology, and raises questions as to whether the relationships between higher level CGE clades can be resolved purely through phylogenetic reconstruction. (Philippe et al. 2011). Comparable difficulties have been reported in studies of other RVT-based phylogenies, where deep branching relationships have similarly remained unresolved despite extensive sampling (Edgar 2022).

### Incorrect classification results in incorrect generalisations at family level and higher

Describing structural or functional features of transposable elements is non-trivial, particularly when such descriptions are intended to be generalisable beyond individual instances. Feature statements such as “LINEs typically encode an RVT domain” or “ORF1 of CR1s contain an esterase” are only meaningful when they can be consistently applied to a biologically valid unit.

The conventional ranks of family and superfamily within LINEs are notoriously unstable (Storer et al. 2021). These ranks nevertheless anchor comparative inference, specifying the scale at which generalisations are intended and establishing an evolutionary context for traits to be interpreted. Put plainly: generalisation is only as strong as the taxonomic framework that underlies it. If the unit to which a trait is attributed is internally heterogeneous or incorrectly assigned, then the trait risks being mis-generalised, misinterpreted, or falsely assumed to be evolutionarily conserved. While this does not preclude classification using currently accepted methods, it does call for such assignments to be applied with appropriate scrutiny given their known limitations.

## Conclusions

This study presents the first comprehensive survey of CR1-group elements across chordates, providing an evolutionary framework that complements and extends current knowledge of CGE diversity and history. Our analyses suggest that several major CGE lineages found in chordates are ancient with deep origins shared across broad chordate groups, and identify additional divergent elements that highlight the extent of unsampled TE diversity. We further show that the persistence and loss of CGEs is not uniform, with the frequency of extinction varying between different CGE clades. We also observed cases in which CGEs within existing databases are assigned classifications that do not align with phylogenetic results, raising concerns that such inconsistencies may propagate systematic errors in future TE classifications. Overall, these results establish a baseline for interpreting CGE evolution and provide a foundation for future studies on the diversification, genomic dynamics, and long-term persistence of these retrotransposons in chordates.

## Methods

### De Novo TE identification

The TE extraction pipeline (https://github.com/alexanderstuart/CR1-Group_Elements_in_Chordates/tree/main) was run in a High-Performance Computing (HPC) environment managed by SLURM using Nextflow scripts. Genomes were downloaded from NCBI using the datasets command-line tool v.15.29.0 (O’Leary et al. 2024). The larger (40Gb) genome of the lungfish was split using the tool seqkit v2.6.1 (Shen et al. 2016), to make the task computationally easier. Previously described and curated CGEs (as of 25.01.24) were downloaded from Repbase (Bao et al. 2015) and Dfam (Storer et al. 2021). Regions within the genomes corresponding to the RVT domain of known CGEs were identified using blastn v2.15.0 (Camacho et al. 2009). The resulting matched seed regions were clustered at 85% sequence identity using cd-hit-est v4.8.1 (Fu et al. 2012) to reduce redundancy; clusters containing rare (fewer than ten) sequences were discarded. RepeatMasker v4.1.5 (Chen 2004) was subsequently used to remove false positive and chimeric sequences. The filtered sequences were then extended using MCHelper v1.6.6 (Orozco-Arias et al. 2024) to automatically extend the seed regions. The output sequences from MCHelper were subsequently inspected using TE-Aid v.1.1-dev (Goubert et al. 2022) and AliView v1.28 (Larsson 2014) to remove false positives. An additional round of extension was performed on sequences that appeared to be prematurely truncated at the 5’ end, which we found was frequently the case in TE families output by MCHelper.

### Phylogenetic Analyses

The consensus sequences generated from our TE extraction pipeline were used to extract largely intact (>80% sequence length, 80% shared sequence identity) representative sequences from the target genomes using blastn (version 2.15.0.) TE consensus sequences with at least 5 copies meeting this threshold were processed further. The RVTs from these genomic copies were extracted using getORF from EMBOSS v6.6.0 (Rice et al. 2000) and hmmsearch from HMMER v3.4 (Potter et al. 2018), searching against the RVT domain included in Pfam database release 37.0 (Mistry et al. 2021), and filtered for minimum length of 220 amino acids. These sequences were aligned using MAFFT v7.525 (Katoh et al. 2002) and a consensus sequence was generated using cons from EMBOSS v6.6.0: these consensus sequences were subsequently clustered at 60% identity using mmseqs2 v16.747c6 (Steinegger & Söding 2017), with the centroid sequence of each cluster selected for further analysis. These newly generated RVT consensus sequences, along with database derived RVT sequences were clustered at 60% identity using mmseqs2, resulting in a reduced dataset of 628 sequences. If a de novo sequence from the same Order as the centroid was included in the cluster, it was given priority over database sequences.

A BEAST XML file was generated using BEAUti v2.7.8 (Bouckaert et al. 2019), using a WAG substitution model and a lognormal relaxed clock model. Ten MCMC analyses were carried out with a chain length of 150,000,000, sampling every 1000 generations in BEAST v2.7.7 (Bouckaert et al. 2019). In TreeAnnotator v2.7.7, 50% of the first trees were discarded and, upon confirming divergence had been achieved, were combined using LogCombiner v2.7.7. TreeAnnotator was then used to generate a Maximum Clade Credibility tree.

The RVT domains of CGE clades identified in the prior Bayesian were realigned using MAFFT v7.525 (Katoh et al. 2002). Maximum likelihood trees for the RVT domains of CGE clades were inferred using IQ-TREE v3.0.1 (Minh et al. 2020), with the optimal model determined by Model-Finder (Kalyaanamoorthy et al. 2017) (Q.pfam+F+R10). iTOL (Letunic & Bork 2024) was used to manage and annotate the generated trees.

### Estimating synonymous and non-synonymous mutation rates of TEs using BUSCO

To estimate the expected level of molecular divergence of each species under vertical inheritance, we calculated synonymous distance (Ks) using single-copy orthologs generated by BUSCO v5.82 (Simão et al. 2015). We adapted scripts from Zhang et al. (2020) and Muller et al. (2024) for our study. In short, we first extracted single-copy orthologs for all species containing CGEs using BUSCO. Then we used MMseqs2 easy-rbh to perform similarity searches to find the best reciprocal hit within the single-copy orthologs (Steinegger & Söding 2017). The corresponding nucleotide and protein regions from these BUSCOs were extracted and used to calculate Ks scores using Li’s method in the seqinr R package. Saturated Ks values (>9) and alignments smaller than 100 aa were filtered, and then the median Ks value was calculated per pair of species. For horizontal transfer, we looked within the 0.5% quantiles of the Ks distribution and the time of divergence.

### Identifying potential HTT candidates

To decrease the workload and to conservatively identify HTT candidates, we implemented several filters. First, we used the reverse transcriptase domains of our identified CGEs (filtered to keep sequences > 200 amino acids). We used Clustal Omega v2.1 to generate a percentage identity matrix based on the amino acid sequence of our dataset (Simão et al. 2015; Sievers et al. 2011). This resulted in a pairwise comparison of each TE based on sequence homology. Then, we filtered to keep only TEs that had at least 75% sequence homology. With our filtered dataset of TEs, we then retrieved the corresponding nucleotide sequence and used it with the amino acid sequences to generate codon alignments using PAL2NAL v14 (Suyama et al. 2006). *Ks* values for each TE-TE pair were estimated using Li’s method in the seqinr R package. If the *Ks* values of the TE-TE pair were smaller than the 0.5% quantile of the BUSCO distribution of their corresponding genomes, we considered this to be a potential HTT event (Supplementary Table 4). We then compared these results with the CR1 phylogeny to determine if the TE pairs were incongruent with the expected host phylogeny, providing further validation. Overall workflow for HTT identification is shown in Supplementary Figure 4.

### Domain annotation

All CGE elements characterised *de novo* in this study were subjected to protein domain annotation. ORFs were predicted with getORF from EMBOSS v6.6.0 (Rice et al. 2000). Predicted amino acid sequences were queried against the Pfam database release 37.0 (1996) using hmmsearch HMMER v3.4 (Potter et al. 2018), and filtered for E-value ≤ 0.05. Domains of interest were manually inspected for HMM profile coverage and sequence complexity, to remove artefactual and low-complexity matches.

### Clustering-Based Inference of Esterase ORF1 Lineage Association

To determine whether individual RVT sequences were affiliated with CGE lineages carrying an esterase-type ORF1, we used sequence clustering to identify patterns of close and distant association based on the similarity of reverse transcriptase domains. Reverse transcriptase (RVT) domains from all sequences were extracted and clustered with MMseqs2 (Steinegger & Söding 2017) at pairwise sequence identity thresholds of 80% and 60%. Cluster membership relative to sequences annotated with an esterase domain in ORF1 was used to assign esterase-association status. A sequence was classified as belonging to a lineage with esterase ORF1 if it was clustered at 80% identity with at least one esterase-annotated sequence. A sequence was classified as distantly related to an esterase-ORF1 lineage if it did not co-cluster at 80% identity with any esterase-annotated sequence, but did co-cluster with such a sequence at 60% identity. A sequence was classified as lacking esterase ORF1 if it did not co-cluster with any esterase-annotated sequence at the 60% identity threshold.

### Comparing patristic distance with pairwise sequence similarity

An all-vs-all similarity search was performed on the RVT sequences of CGEs included in the BEAST analysis using mmseqs search (MMseqs2) (Steinegger & Söding 2017). For each query sequence, the top-scoring hit was defined as the non-self match with the highest bitscore. Patristic distances for the corresponding sequence pairs were computed from the BEAST tree in R using cophenetic.phylo (ape v5.8.1).

