## Supplementary_Figures for "Ancient persistence and newfound diversity of CR1-group retrotransposons across chordates"

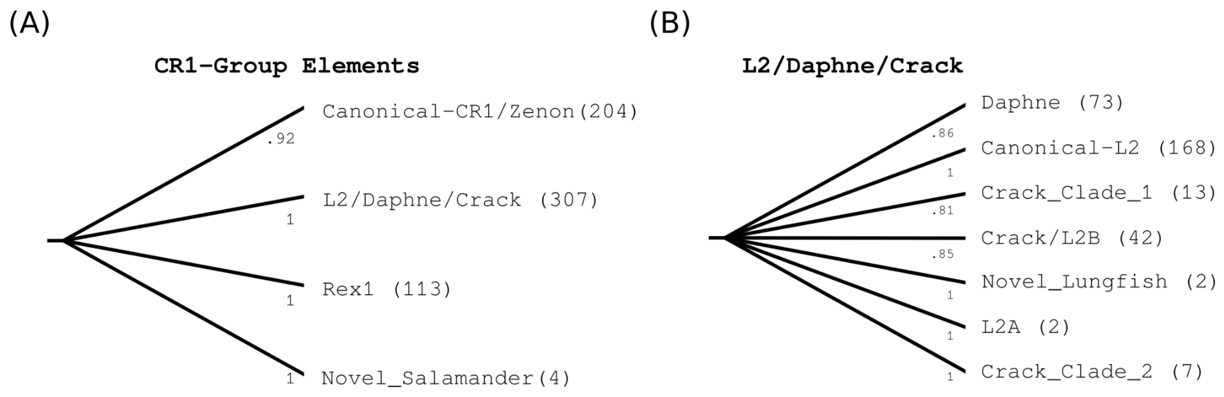


**Supplementary Figure 1: Bayesian phylogeny of CGEs.** The numbers in parentheses indicate the total number of sequences collapsed into that clade. Clades are named based on the most frequent TE classifications among the sequences they contain. (A) Tip-collapsed representation of the Bayesian phylogeny inferred from the conserved RVT domain of the 628 CGEs analysed in this study. The backbone of the tree is unresolved, forming a polytomy due to low posterior probability values at deep nodes. This polytomy comprises four major clades, each of which is well supported by posterior probability values between 0.92 and 1. (B) Tip collapsed view of the L2/Daphne/Crack clade from Panel (A), showing the next level of high confidence subclades. These subclades form an unresolved polytomy, with each subclade well supported by posterior probabilities between 0.81 and 1.

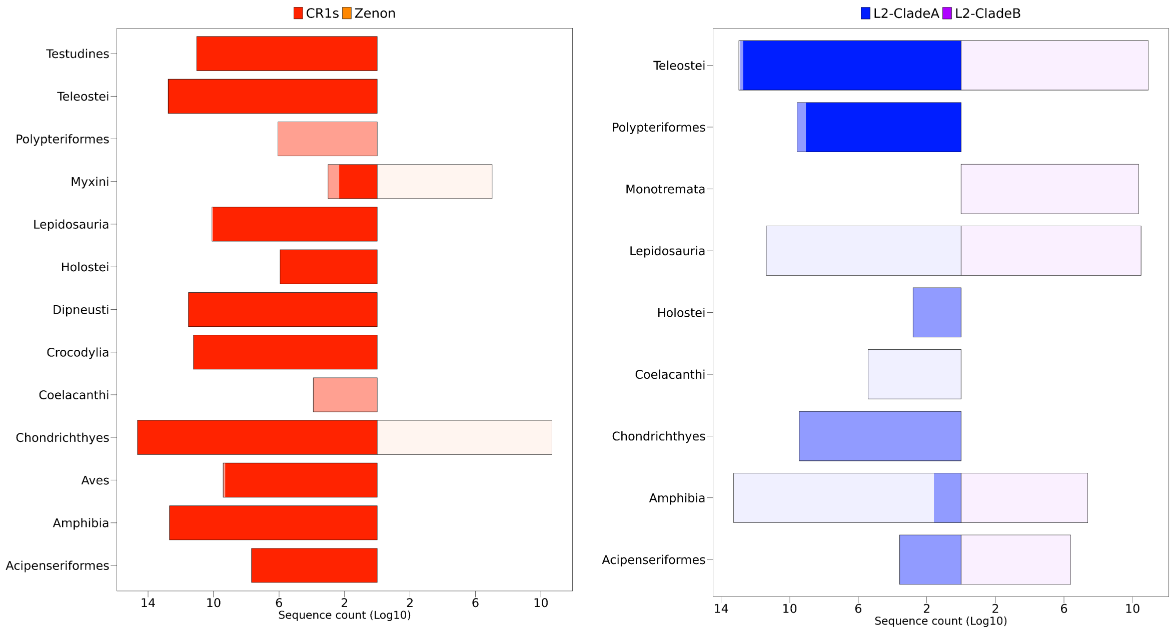


**Supplementary Figure 2: Distribution of Esterase ORF1 across different lineages of CR1, Zenon, L2 clade A and B.** the different shades represents number of sequences that are: **(solid)** clustered with esterase carrying CGE at 80% ID; **(medium shade)** cluster with clustered with esterase carrying CGE at 60% ID; **(low shade)** does not cluster with esterase carrying CGE.

Template alignment | Template 3D Structure | PDBe

1. **2YKO_C LINE-1 ORF1P; RNA-BINDING PROTEIN, GENOME EVOLUTION, NUCLEIC ACID CHAPERONE, RNP, COILED-COIL, RNA BINDING PROTEIN; HET: MSE; 2.1A {HOMO SAPIENS}**

Probability: 98.91%, E-value: 8.8e-7, Score: 64.81, Aligned cols: 155, Identities: 14%, Similarity: 0.221, Template Neff: 11.4

```
Q ss_pred              HHHhhhHHHHHHHHHHHHHHhcceEEeeeeecccchhcccccccCccHHhHHcHHHHHHHHHHhcCCcc---Cccccccee
Q GouWil_Daphne-   71  AEKIKTLDARVEDLERKEREKDVIITGLKIRPRSYASATRQTEDQTIEEEKSIEEQVIDYLDSKGNEV---NPDNINYFQ   147 (244)
Q Consensus        71  aekiktldarvedlerkerekdviitglkirprsyasatrqtedqtieeeksieeqvidyldskgnev---npdninyfq   147 (244)
                       ..++..+..++++++...+...++|.|+.-.+..          ...    .+...+...+...-..+   .+..+....
T Consensus        33  ~~~~~~l~~~~~~~e~~~r~~ni~i~gi~e~~~~----------~~~----~~~~~~~~~l~~~~~~~~~~~~~~~i~~~~    98 (233)
T 2YKO_C         33  EKRIKRNEQSLQEIWDYVKRPNLRLIGVPESDVE----------NGT----KLENTLQDIIQENFPNLARQANVQIQEIQ    98 (233)
T ss_dssp             HHHHHHHHHHHHHHHHHHTTTEEEEESCCCCSSC----------CHH----HHHHHHHHHHHHHCGGGGSSSCCCEEEEE
T ss_pred             HHHHHHHHHHHHHHHHHhhcCCEEEeCCCcccCC----------Cch----hHHHHHHHHHHHhCchhHHhcCcceeEee

Q ss_pred              cccC-----CCCCCcceeEEechhHHHHHHHhhhh-----hccEEEeecccccccchh---HHHHHHHHHcCCeeeeccc
Q GouWil_Daphne-  148  LLPK-----RKDTRDVKITFTNVKYKAEIMKQGRK-----LAGTMVFMNENLTKQNAS---IAWKARQLKKGGKIVKNWT   214 (244)
Q Consensus       148  llpk-----rkdtrdvkitftnvkykaeimkqgrk-----lagtmvfmnenltkqnas---iawkarqlkkggkivknwt   214 (244)
                       .+++     ....+.+.+.|.+...+..+++..++     +.|+-+++++.++.....   .-.+.++..+...+...|+
T Consensus        99  r~~~~~~~~~~~~~r~i~v~f~~~~~~~~~~~l~~~~~~~~~~~~~~~~~~~~~~i~i~~~l~~~~~~~~~~~~~~~~~~~~~~~~~~~~~~~~   178 (233)
T 2YKO_C         99  RTPQRYSSRRATPRHIIVRFTKVEMKEKMLRAAREKGRVTLKGKPIRLTVDLSAETLQARREWGPIFNILKEKNFQPRIS   178 (233)
T ss_dssp             ECC----------CCEEEEEESCHHHHHHHHHHHHHCCCEETTEECEEEECCCHHHHHHHHHHTHHHHHHHHTTCCCEEE
T ss_pred             cCCCCCCCCCCCCCcEEEEeCCHHHHHHHHHHHHHHhCCceeCCeEEEEeCCCCHHHHHHHhHHHHHHHHHHCCCCcccc

Q ss_pred              CCceEEecCCCCcCCceeeeeehhHHhhc
Q GouWil_Daphne-  215  RGCRIYITPPGGENGKPVLVQRMEDLEKY   243 (244)
Q Consensus       215  rgcriyitppggengkpvlvqrmedleky   243 (244)
                       ..+++++..    +|++..+...++++++
T Consensus       179  ~~~~i~~~~----~~~~~~~~~~~~~~~~   203 (233)
T 2YKO_C        179  YPAKLSFIS----EGEIKYFIDKQMLRDF   203 (233)
T ss_dssp             TTTEEEEEE----TTEEEEESSHHHHHHH
T ss_pred             CCcEEEEEE----CCEEEEECCHHHHHHH
```

**Supplementary Figure 3:** HHpred results showing remote homology of a GouWI Daphne ORF1 to Transposase_22

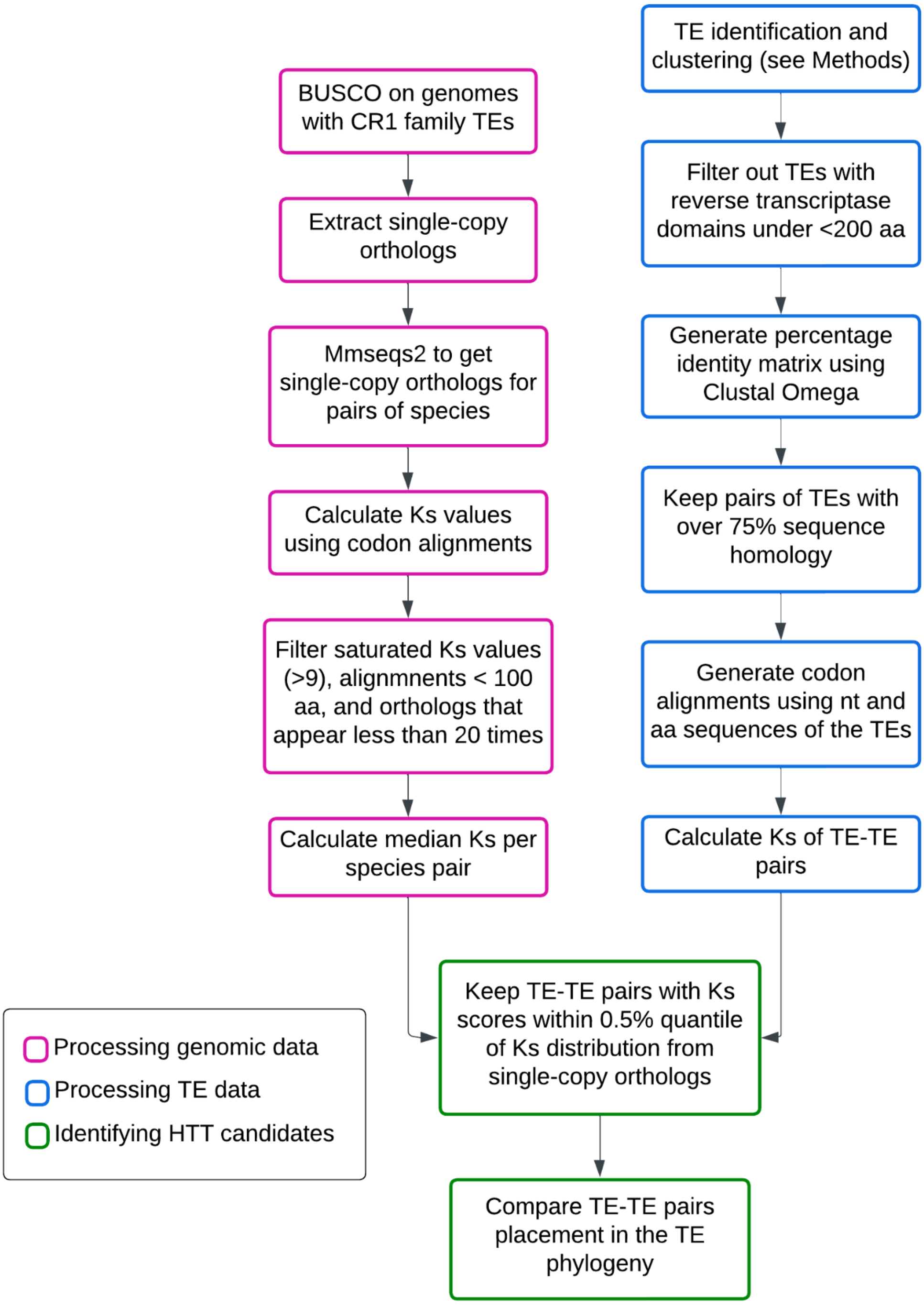


**Supplementary Figure 4: Overview of the methods to identify HTT candidates**. The first processes include getting the BUSCOs for each genome in the study and extracting the single-copy orthologs to calculate median Ks values. The second is to filter TE candidates for HTT and calculate the Ks values of TE-TE pairs. Then use the median Ks values to determine whether there is a likely HTT event. 'HTT' = horizontal transfer of TEs, 'aa' = amino acids, 'nt' = nucleotide, 'Ks' =
